# Genetic Diversity Study of *Fusarium culmorum*: Causal agent of wheat crown rot in Iraq

**DOI:** 10.1101/341909

**Authors:** O. N. Matny, S. A. Shamsallah, M. Haas

## Abstract

Fusarium crown rot (FCR), caused by *Fusarium culmorum* (Wm.G.Sm) Sacc., is an important disease of wheat both in Iraq and other regions of wheat production worldwide. Changes in environmental conditions and cultural practices such as crop rotation generate stress on pathogen populations leading to the evolution of new strains that can tolerate more stressful environments. This study aims to investigate the genetic diversity among isolates of *F. culmorum* in Iraq. Twenty-nine samples were collected from different regions of wheat cultivation in Iraq to investigate the pathogenicity and genetic diversity of *F. culmorum* using the REP-PCR technique. Among the twenty-nine isolates of *F. culmorum* examined for pathogenicity, 96% were pathogenic to wheat at the seedling stage. The most aggressive isolate, from Baghdad, was IF 0021 at 0.890 on the FCR severity index. Three primer sets were used to assess the genotypic diversity via REP, ERIC and BOX elements. The amplicon sizes ranged from 200-800 bp for BOX-ERIC2, 110-1100 bp for ERIC-ERIC2 and 200-1300 bp for REP. In total, 410 markers were polymorphic, including 106 for BOX, 175 for ERIC and 129 for the REP. Genetic similarity was calculated by comparing markers according to minimum variance (Squared Euclidean). Clustering analysis generated two major groups, group 1 with two subgroup 1a and 1b with 5 and 12 isolates respectively, and group 2 with two subgroups 2a and 2b with 3 and 9 isolates respectively. This is the first study in this field that has been reported in Iraq.

## INTRODUCTION

*Fusarium culmorum* (Wm.G.Sm) Sacc., a fungal plant pathogen with a wide host range and is the causal agent of several diseases on these plants. On wheat, *F. culmorum* causes two important diseases that can cause serious economic losses on wheat: head blight and crown rot (Burgess et al. 2001; Chakraborty et al. 2006). Reliable estimates for yield loss due to Fusarium crown rot (FCR) in Iraq are not available, but where data are available, FCR can be devastating. For example, FCR can reduce yields of winter wheat production in the Pacific Northwest region of the USA by up to 61% (Smiley et al. 2005). FCR also affects grain quality through the production of mycotoxins such as DOV, NIV, ZEN and T2-toxin which can be harmful to human, and livestock health (Pestka and Smolinski 2005; Blandino et al. 2012).

Over the past 5 years, FCR re-emerged as an economically important disease in Iraq, causing significant yield losses to the wheat crop (Matny et al. 2012). A few studies on FCR have been carried out to understand the genetic diversity present in *F. culmorum* populations and to understand why this disease has re-emerged. Drought conditions in Iraq from 2011 to 2016 are likely a contributing factor to the spread of *F. culmorum* in wheat fields since many studies have shown that dry environments are favorable for *F. culmorum* growth and reproduction (Scherm et al. 2013; Balmas et al. 2006). FCR has also been reported in other Middle Eastern countries such as Turkey (Tunali et al. 2006; Emre et al. 2016), Iran (Hajieghrar 2009; Eslahi 2012) and Syria (Khalifeh et al. 2009).

Genetic diversity analyses of microorganisms have demonstrated that pathogen diversity depends on global environmental changes and shifts in agro-ecological systems (Saharan and Naef 2008; Gurel et al. 2010). *F. culmorum* isolates show high levels of phenotypic and genotypic variability in culture, including colony morphology, pigmentation and sporulation (Puhalla 1981; Kollers et al. 2013; Miedaner et al. 2013). In addition, variation in aggressiveness and mycotoxin production have been found among various isolates that were collected from different geographic locations (Gang et al. 1998; Berna et al. 2012; Winter et al. 2013; Fang et al. 2015).

There are many methods and techniques used for studying the genetic diversity of microorganisms, including Repetitive Polymerase Chain Reaction (REP-PCR), also known as Repetitive DNA-based fingerprinting. The amplification of prokaryotic genomic sequences between the repetitive elements include: Repetitive Extragenic Palindromic (REP) sequences, Entero-bacterial Repetitive Intergenic Consensus (ERIC) sequences and BOX elements. REP-PCR applications are in widespread use among studies of plant pathogenic bacteria, but among eukaryotic microbes, have only been tested in *F. oxysporum* (Edel et al. 1995). The principle aim of this study is to characterize the genetic diversity among *F. culmorum* isolates collected in Iraq through REP-PCR and associate the results with their geographic distribution and pathogenicity toward wheat at the seedling stage.

## MATERIALS AND METHODS

### Plant material and fungal isolation

Wheat plants exhibiting FCR symptoms were collected from seven provinces representing different agricultural zones in Iraq. All samples were collected in paper envelopes and necessary data (sample number, place and date of collection and host cultivar name) were recorded. The samples were brought to the laboratory and kept in a well-ventilated area at room temperature (25±3°C and 30% humidity) until the samples could be processed. The crown of wheat plants were cut into 0.5-1.0 cm segments, and treated with a 10% sodium hypochlorite (bleach) solution (diluted from commercially-available concentrated bleach) for 2 min, washed with sterile water and dried with filter paper. All samples were placed into 9 cm petri dishes containing potato dextrose agar (PDA). Fifty milligrams of Agrimycin-343 was added to the medium after autoclaving. Plates were incubated at 25°C±2 for 5 days and then a single *Fusarium* spp. spore was picked up (under a microscope at 400X) using a needle according to colony and spore characterization methodologies (Booth 1971; John and Brett 2006) and placed into a new petri dish containing PDA for use in pathogenicity assays.

### Pathogenicity assay

For the pathogenicity assays, *F. culmorum* isolates were grown on autoclaved millet seed. First, one kilogram of millet seed was soaked in water for 12 h, then the water was drained, and several 250 ml flasks were filled with 50 g each of this millet seed and autoclaved at 121°C and1.5 kg cm^-1^ pressure for 20 min. One disc (0.5 cm) of a 7-day old *F*. *culmorum* colony was placed into each flask and incubated at 25±2°C for 14 days. Pathogenicity tests were then performed on wheat seedlings. A 1:1 mixture of sterile soil and peat moss was autoclaved at 121 °C and 1.5 kg cm^-1^ pressure for 1h. This process was repeated on two separate days. Pots (5 × 10 cm) used in the greenhouse experiments were filled with the mixture soil and 5 g of *F. culmorum* inoculum was added to each pot in the top 5 cm surface layer of the soil. All pots were watered and placed in the greenhouse for 2 days 25±2. Three seeds of *Triticum aestivum* L. cv. Abu-Ghreeb1 (a commonly used cultivar used in Iraq) were sown in each pot, and each treatment repeated three times. Plants were irrigated with sterilized water as needed. After 35 days, crown rot symptoms and stem discoloration characteristic of FCR were apparent on the inoculated plants. FCR Severity was calculated by measuring the length of discoloration relative to seedling height. The FCR severity index was obtained by multiplying this ratio by the number of leaf-sheath layers with necrosis. The FCR index was calculated according to the following formula: (length of stem discoloration/seedling height) × (number of leaf sheath layers with necrosis) (Mitter et al. 2006).

### Fungal growth and DNA extraction

Twenty-nine *Fusarium* ssp. isolates were grown on PDA media in 9 cm petri dish for 7 days at 25±2°C. The REDExtract-N-AMP™ Plant PCR kit (Sigma-Aldrich, St. Louis, MO, USA) was used for DNA extraction according to manufacturer’s instructions. Briefly the hyphal tip of the mycelium was harvested with a sterilized needle and placed in 0.2 ml collection tubes, to which 50 µl extraction solution was added, followed by incubation at 95°C for 10 min. The DNA concentrations and quality were measured to ensure quality using a Nanodrop 2000 spectrophotometer (Thermo Fisher Scientific, USA), qualitative analyses of DNA were carried out via agarose gel electrophoresis.

### Amplification of fungal DNA

Species identification of *Fusarium* spp. isolates was determined by amplifying and sequencing the Translation Elongation Factor 1 alpha (TEF-1α) gene. Forward (EF1) 5’-ATGGGTAAGGA(A/G)GACAAGAC-3’ and reverse (EF2) 5’-GGA(G/A)GTACCAGT(G/C)ATCATG-3’ (O’Donnell et al. 2000) primers were used to amplify the TEF1 α gene. The PCR reaction solution was prepared at total volume of 20 µl. The PCR conditions were as follows: an initial denaturation at 95°C for 5 min followed by 35 cycles of denaturation at 94°C for 50 s, annealing at 53°C for 50 s, extension at 72°C for 1 min, and a final extension at 72°C for 7 min. Amplification products were visualized on 1.0% agarose gel stained with SYBR^TM^ safe DNA gel stain in 1X TAE (Invitrogen™, Carlsbad, CA, USA). Amplification of a product approximately 700 bp long was generated by PCR from the DNA template.

### DNA sequencing

The PCR products of TEF-1α were prepared for sequencing by cleaning up with the QiAquick^®^ PCR purification kit (city? Bogota?, Colombia). Sequencing was carried out commercially (ACGT, Inc., Chicago, USA). The sequencing chromatograms were read and aligned using MEGA6 software and the sequences were compared with those in GenBank (http://www.ncbi.nlm.nih.gov/) for the TEF-1α gene using the Basic Local Alignment Search Tool (BLAST). All sequences of the isolates were sent to GenBank to obtain accession numbers.

### Genetic diversity study

*F. culmorum* isolates collected from different regions of Iraq were genotyped using three primer pairs designed to amplify multiple regions of the genome simultaneously: (1) REP1R-Dt (5′-III NCGNCATCNGGC-3′) and REP-2G (5′-GCGGCTTATCGG GCCTAC-3′) for REP; (2) ERIC 1 (5′-ATG TAAGCTCCTGGGGATTCAC-3′) and ERIC 2 (5′-AAGTAAGTGACTGG GGTGAGCG-3′) for ERIC, and (3) BOX-A1R (5′-CTACGGCAAG GCGACGCTGACG-3′) (Versalovic et al. 1991) for BOX-ERIC2. The PCR conditions were: an initial denaturation at 94°C for 5 min, followed by 40 cycles of 94°C for 1 min, an annealing step of 50°C (BOX and ERIC) or 37°C (REP) for 1 min, and an extension at 72°C for 2 min, followed by a final extension at 72°C for 15 min. Amplification products were visualized on 1.5% agarose gels with SYBR safe DNA gel stain in 1X TAE (Invitrogen™). In total, 410 potential markers were generated by this method for genotyping. REP-PCR markers were evaluated together in pair-wise comparisons. Single and shared fragments were analyzed using by Multivariate Statistical Package (MVSP) 3.22 program, the similarity was calculated according to minimum variance (Squared Euclidean) (Kovach 2001).

## RESULTS

Pathogenicity tests for *F. culmorum* on wheat seedlings demonstrated that the collected isolates vary in their pathogenicity toward wheat cultivar Abu-Grheeb1, ranging from 0.001 to 0.890 on the FCR severity index; however, some isolates (IF 0003, IF 0013, IF 0024) were non-pathogenic, with a score of 0.00 on the FCR severity index. The isolates that resulted in the highest FCR severity index scores were: IF 0021, IF 0028, IF 0045, IF 0046, IF 0015, and IF 0005. Isolates IF 0021, IF 0028, IF 0045 and IF 0046 originate from Baghdad while IF 0015 and IF 0005 come from Anbar and Diyala provinces, respectively (Table 1).

**Table 1.** *Fusarium culmorum* accessions used in this study including loci information, TEF-1α gene test and disease severity index.

| No. | Accession<br>No. | Isolate<br>No. | Species | Location | TEF-1α<br>gene | FCR<br>severity<br>index |
| --- | --- | --- | --- | --- | --- | --- |
| 1 | KY205745 | IF 0003 | *F. culmorum* | Karbala | + | 0.00 |
| 2 | KY205746 | IF 0004 | *F. culmorum* | Karbala | + | 0.030 |
| 3 | KY190104 | IF 0005 | *F. culmorum* | Diyala | + | 0.286 |
| 4 | KY190106 | IF 0006 | *F. culmorum* | Diyala | + | 0.001 |
| 5 | KY190111 | IF 0007 | *F. culmorum* | Diyala | + | 0.074 |
| 6 | KY190107 | IF 0008 | *F. culmorum* | Diyala | + | 0.002 |
| 7 | KY190127 | IF 0009 | *F. culmorum* | Kirkuk | + | 0.098 |
| 8 | KY190118 | IF 0013 | *F. culmorum* | Anbar | + | 0.00 |
| 9 | KY190123 | IF 0014 | *F. culmorum* | Anbar | + | 0.011 |
| 10 | KY190116 | IF 0015 | *F. culmorum* | Anbar | + | 0.309 |
| 11 | KY190108 | IF 0017 | *F. culmorum* | Najaf | + | 0.022 |
| 12 | KY205747 | IF 0021 | *F. culmorum* | Baghdad | + | 0.890 |
| 13 | KY190121 | IF 0022 | *F. culmorum* | Baghdad | + | 0.294 |
| 14 | KY190126 | IF 0024 | *F. culmorum* | Diyala | + | 0.00 |
| 15 | KY190122 | IF 0026 | *F. culmorum* | Anbar | + | 0.050 |
| 16 | KY190112 | IF 0028 | *F. culmorum* | Baghdad | + | 0.543 |
| 17 | KY205748 | IF 0029 | *F. culmorum* | Baghdad | + | 0.004 |
| 18 | KY190109 | IF 0030 | *F. culmorum* | Kirkuk | + | 0.001 |
| 19 | KY190113 | IF 0031 | *F. culmorum* | Kirkuk | + | 0.030 |
| 20 | KY190114 | IF 0032 | *F. culmorum* | Kirkuk | + | 0.002 |
| 21 | KY190117 | IF 0033 | *F. culmorum* | Kirkuk | + | 0.004 |
| 22 | KY190124 | IF 0040 | *F. culmorum* | Babylon | + | 0.076 |
| 23 | KY190110 | IF 0041 | *F. culmorum* | Babylon | + | 0.005 |
| 24 | KY190119 | IF 0042 | *F. culmorum* | Babylon | + | 0.001 |
| 25 | KY205749 | IF 0044 | *F. culmorum* | Diyala | + | 0.019 |
| 26 | KY190105 | IF 0045 | *F. culmorum* | Baghdad | + | 0.350 |
| 27 | KY190125 | IF 0046 | *F. culmorum* | Baghdad | + | 0.200 |
| 28 | KY190120 | IF 0047 | *F. culmorum* | Baghdad | + | 0.008 |
| 29 | KY190115 | IF 0052 | *F. culmorum* | Anbar | + | 0.010 |

The results of species-specific identification using TEF-1α demonstrated that all isolates used in this study are in fact *F. culmorum*.

The genetic diversity study showed monomorphic and polymorphic bands pattern. The annealing temperature for the BOX, ERIC and REP primers used in this study were different from the output provided by Gurel et al. (2010). In this study, we found the optimum annealing temperature for the primers was 50°C for BOX and ERIC, and 37°C for REP. The PCR bands for the final amplification products were between 200-800 bp for BOX-ERIC2, 110-1100 bp for ERIC-ERIC2, and 200-1300 bp for REP. A total of 410 polymorphic markers were identified in this study for the *F. culmorum* isolates, including 106 for BOX, ERIC for 175 and 129 for the REP-PCR. These markers were used to evaluation the minimum variance between the strains (Fig. 2).

Minimum variance cluster analysis was used to detect the variance between the *F. culmorum* isolates (fig.1). The dendrogram illustrates separated the isolates into major groups in this present study. Group 1 includes 17 isolates and may be divided into two sub-groups (1A and 1B) consisting of 5 and 11 isolates, respectively. All isolates in this group were collected from northern and central sites in Iraq. Group 2 consists of 12 isolates and may also be divided into two subgroups (2A and 2B) consisting of 3 and 9 isolates, respectively. Group 2A isolates originate from central Iraq while group 2B isolates come from northern, central, and southern Iraq.

**Figure 1.**
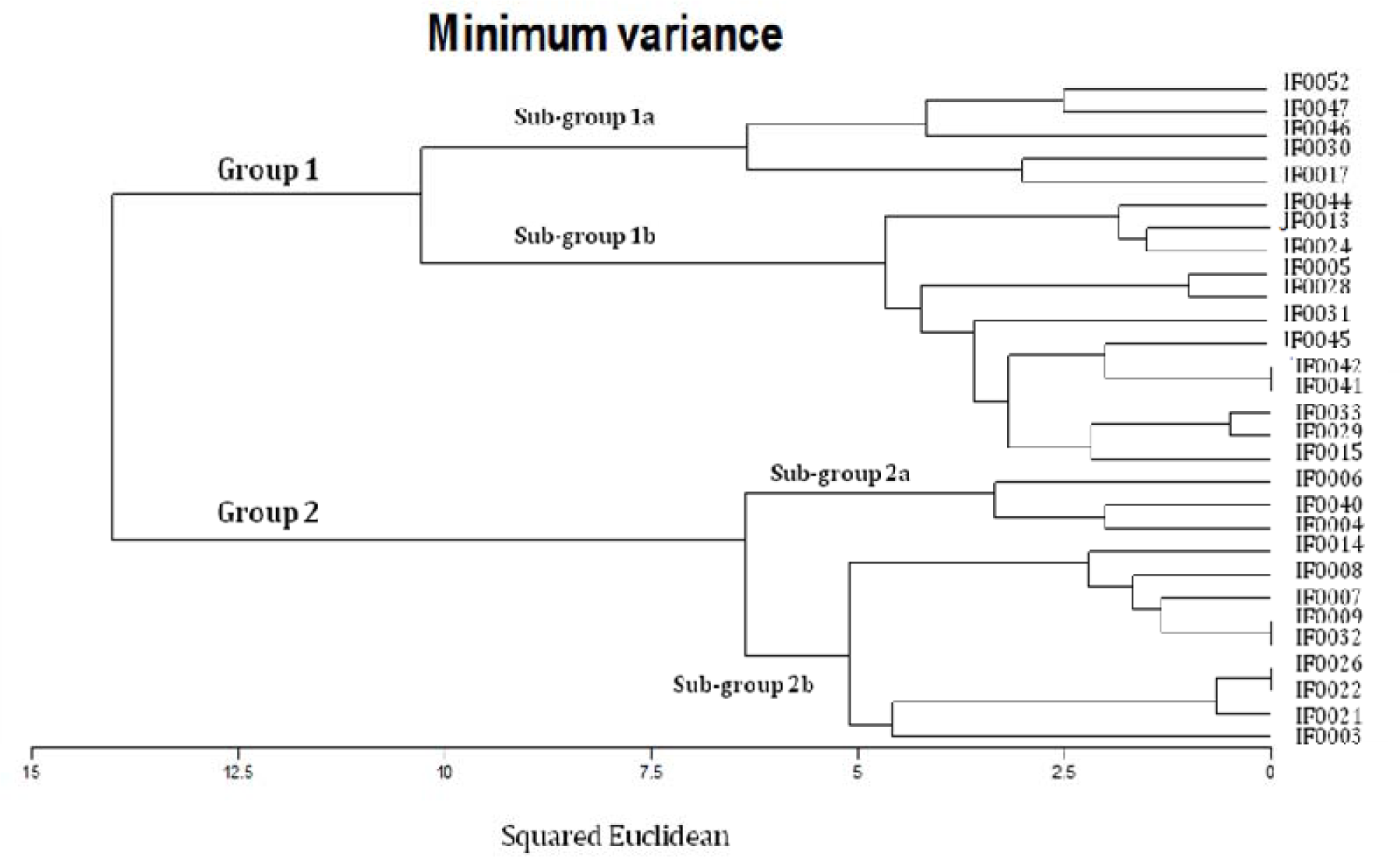
Dendrogram of 29 *Fusarium culmorum* isolates generated based on the number of bands and position of appearance for three primers BOX-ERIC2, ERIC-ERIC2 and REP by using Multivariate Statistical Package (MVSP) 3.22 program to show minimum variance (Squared Euclidean).

**Figure 2.**
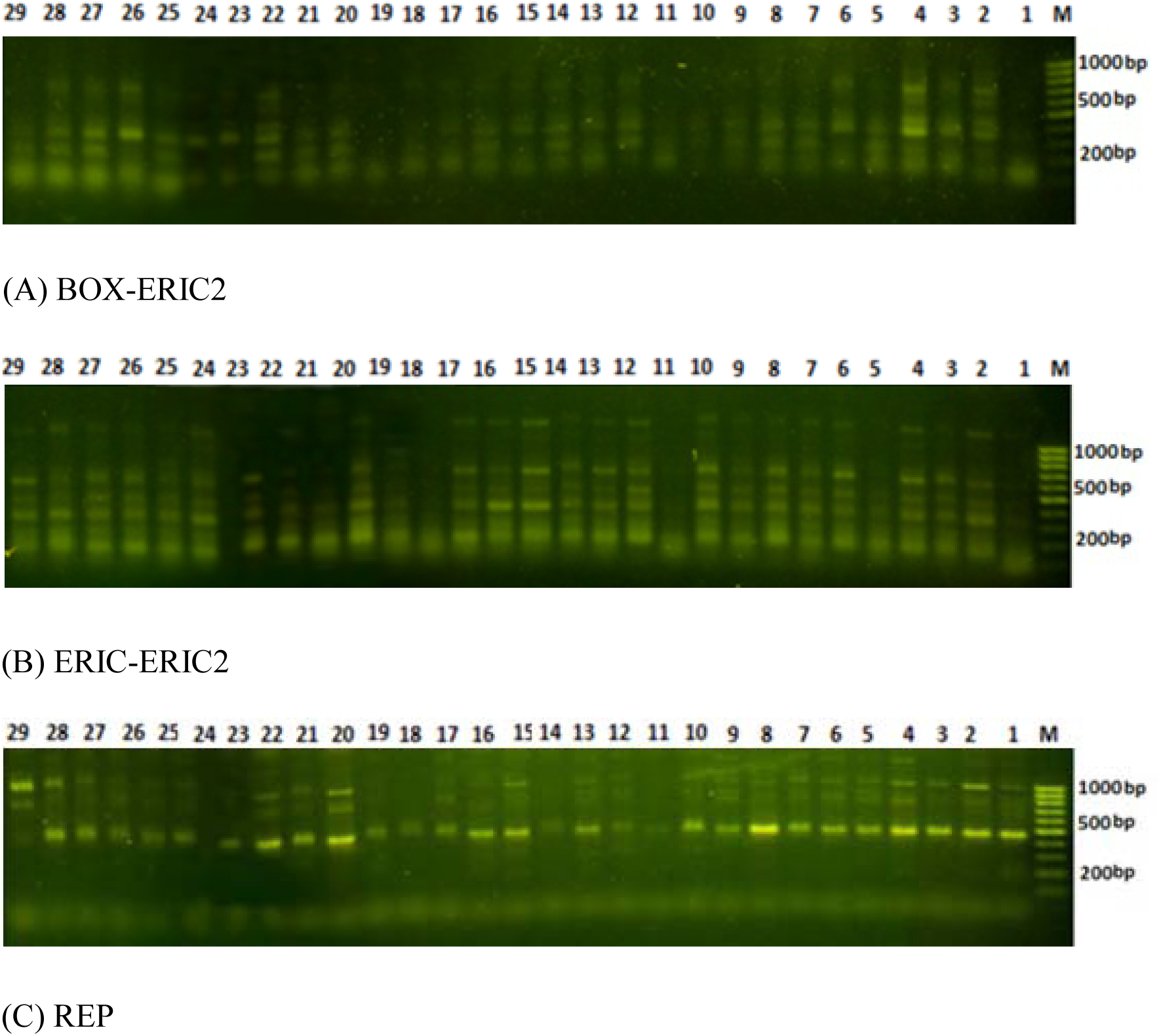
Fingerprint pattern for 29 *F. culmorum* isolates by using three primer pairs: (A) BOX-ERIC2, (B) ERIC-ERIC2 and (C) REP.

Isolates IF0022 and IF0026 share the highest similarity values (100% similar). There is no relationship between the geographic origin of the isolate and its genetic relationship to other isolates (Fig. 1). One of the reasons is because of the use of seed that has not been certified by the Iraqi Ministry of Agriculture and the exchange of seed between farmers across different regions leading to the transfer the pathogen with the seed from one province to another. Also, some farmers obtain their seed from local markets where the seed source is unknow.

## DISCUSSION

Genetic diversity studies of pathogen populations are important for understanding the genetic potential of economically important pathogens to adapt to climate change and the implications for management of diseases caused by these pathogens (McDonald and Linde 2002). *F. culmorum* has also been reported to cause seed-borne diseases of pre- and post-emergence seedling death and is one of the causal species of Fusarium head blight (FHB) (Polley and Turner 1995). In this study, **89.7%** of the *F. culmorum* isolates were pathogenic towards wheat at the seedling stage, while a smaller **10.3%** number of isolates were non-pathogenic.

As the results of our study of the genetic diversity among 29 isolates of *F. culmorum* collected from different regions of Iraq show, there is no relationship between geographic location and genetic similarity of the isolates. We suggest that this means that *F. culmorum* populations have the ability to survive and adapt to different and extreme variations in climate, from the cold area in northern Iraq with temperatures ranging from −5 to 10°C in winter and 35 to 45°C in summer; to central and southern Iraq where temperatures range from around 5 to 15°C during winter to 45 to 55°C in the summer.

Many studies of DNA analysis have been reported to investigate genetic variability and population structure of *F. culmorum* using a variety of molecular markers, such as random amplified polymorphic DNA (RAPD) (De Nijs et al. 1995; Gargouri et al. 2003; Yörük and Albayrak, 2013) and restriction fragment length polymorphism (RFLP) (Nicholson et al 1993; Llorensa et al. 2006). These studies suggest that there is extensive genetic diversity in *F. culmorum* populations. In addition, Mishra et al. (2003) found a high degree of intra-specific polymorphism among *F. culmorum* isolates using inter-simple sequence repeat (ISSR) analysis. Albayrak et al. (2016) also studied the relationship between *Fusarium* isolates according to their species and geographic regions by using ISSR markers. In another study, Bayraktar and Fatma (2010) found that ISSR markers have a high degree of intra- and interspecific polymorphisms among *Fusarium* spp. Finally, Gargouri et al. (2003) used RAPD markers to study the genetic variability and population structure of *Fusarium culmorum* isolated from wheat stem bases.

In addition to the genetic diversity and variation in pathogenicity towards wheat presented in this study, populations of *F. culmorum* are also characterized by high levels of phenotypic variability in culture, such as sporulation, pigmentation, mycotoxin production, and colony morphology (Puhalla 1981). Gang et al. (1998) found a large variation among *F. culmorum* isolates collected from the various geographic areas for aggressiveness, race designation and mycotoxin production.

## CONCLUSION

This is the first report of the genetic diversity of *F. culmorum* populations present in Iraq using the REP-PCR method. Two groups of *F. culmorum* were identified in this study according to minimum variance (Squared Euclidean). This is the first study in this field that has been reported in Iraq. Although the study was limited to 29 isolates due to the difficulty in completing more extensive sampling, this study provides a first glimpse at the genetic diversity and variation in pathogenicity present in *F. culmorum* populations in Iraq.

## ACKNOWLEDGMENTS

This work was carried out and supported by the University of Minnesota, Department of Plant Pathology. Thanks to Dr. Scott Bates and Dr. Zewei Song for all the support and help to complete this paper.

